# White adipose tissue undergoes pathological dysfunction in the TDP-43A315T mouse model of amyotrophic lateral sclerosis (ALS)

**DOI:** 10.1101/2025.07.03.662925

**Authors:** Cristina Benito-Casado, Esther Durán-Mateos, Águeda Ferrer-Donato, Gemma Barroso García, Raúl Domínguez-Rubio, Mónica Povedano, Carmen M. Fernandez-Martos

## Abstract

White adipose tissue (WAT) has a crucial role in maintaining systemic energy homeostasis. Numerous biological pathway studies have highlighted the importance of adipokines in regulating metabolic pathways and contributing to metabolic dysfunction in animal models and patients with ALS. Despite these associations, the specific molecular mechanisms remain poorly understood. Moreover, the direct contribution of WAT to the energy metabolism abnormalities observed in ALS has yet to be clearly defined. The current study sought to identify perturbances in WAT, main source of leptin, during the clinical course of the disease in TDP-43^A315T^ mice using histological, proteomic, and molecular biological techniques. We present the first evidence of a significant histological alteration in WAT prior to the symptomatic stage of the disease in TDP-43^A315T^ mice, providing novel insights into pathological features earlier in the onset of symptoms, and showing WAT as a target organ for ALS. In human ALS cases, we found that circulating leptin levels at the time of diagnosis were lower in the plasma of men with ALS who were overweight or obese and had rapidly progressive ALS, emphasizing the importance of considering sex-specific approaches when analysing adipokines essential for body weight control.

## INTRODUCTION

Amyotrophic lateral sclerosis (ALS) is the third most common neurodegenerative disease worldwide characterized by the selective loss of motor neurons both in the brain and spinal cord (Rowland & Shneider, 2001), leading to paralysis and respiratory failure. ALS is often rapidly progressive, incurable, and fatal. Over 60% of patients die within 3-5 years after diagnosis (Connolly et al., 2015). Although much effort has been made in the past two decades to understand the complexity and heterogeneity of ALS, the development of effective therapies remains elusive due to several challenges. One major obstacle is the limited understanding of the underlying mechanisms driving ALS disease.

Identifying the mechanisms involved in maintaining energy homeostasis in ALS can help to elucidate the growing evidence supporting a strong metabolic component in ALS. Indeed, rapid weight loss is associated with worse disease outcomes in ALS (Janse van Mantgem et al., 2020), and fat mass loss correlates with faster disease progression (Lee et al., 2021). Conversely, higher initial body mass index (BMI) and obesity (Heritier et al., 2015), type 2 diabetes mellitus (Jawaid et al., 2010), and patients with higher triglyceride and cholesterol levels (Dorst et al., 2011) have been associated with longer survival in ALS, indicating fat content and lipid metabolism are important prognostic factors in ALS disease. However, despite these correlations, the underlying mechanisms driving these metabolic derangements remain unclear. Adipose tissue is an endocrine organ and constitute the major lipid store in mammals (Kershaw & Flier, 2004). Two main types of anatomically and physiologically different adipose tissue are present in adults: WAT, which is mainly an energy store (Luo & Liu, 2016); and the brown adipose tissue (BAT), with specialized functions in heat production (thermogenesis) (Li et al., 2019). Moreover, as an endocrine organ, adipose tissue secretes a variety of soluble mediators, with a critical role in the physiological regulation of the metabolism acting on the hypothalamus, a key integrative brain area that regulates neuronal and metabolic circuits involved in energy intake regulation (Goel et al., 2025). While dysfunctions in the hypothalamus have only recently been described (Rosina et al., 2025), studies have shown this brain structure is atrophied in ALS (Tse et al., 2023), even in premorbid stages, and correlates with BMI (Gorges et al., 2017). Altered eating behaviour have been reported in patients with ALS and frontotemporal dementia (FTD) (Ahmed et al., 2016). Indeed, we and others demonstrated increased mRNA expression levels of *agouti-related peptide* (AgRP), while levels of its *antagonist pro-opiomelanocortin* (POMC) were decreased in the hypothalamus in TDP-43^A315T^ mice (Ferrer-Donato et al., 2021) and mutant SOD1^G86R^ mice (Vercruysse et al., 2016), demonstrating of a hypothalamic involvement in ALS. Indeed, there are several adipokines involved in metabolism and regulation of food intake which act within the brain, specifically in the brainstem and hypothalamus, including, but not limited to, leptin, which is primarily produced by WAT in proportion to fat stores (Pico et al., 2022). Nevertheless, although clinical evidence suggests that the distribution and content of adipose tissue, and subsequently leptin levels, are significantly altered in ALS and have been correlated with functional status and survival in ALS (Ferrer- Donato et al., 2021; Ngo, S. T. et al., 2015; Picher-Martel et al., 2023), it is currently unclear whether metabolic changes in ALS are driven by the adipose tissue dysfunction, or whether hypothalamic neuronal dysfunction may contribute to disease processes in ALS.

Notably, metabolic changes are also observed in FTD, which exists on a continuous clinical spectrum with ALS (Chen-Plotkin et al., 2010). Indeed, although the clinical manifestations of ALS and FTD differ (Chen- Plotkin et al., 2010), altered peripheral levels of leptin have been found in patients with FTD (Ahmed et al., 2019), while there is also limited information on the mechanism underlying the association between adiposity and brain metabolism. Furthermore, we have determined complex alterations in metabolic hormones, including leptin, in pathology-rich regions of post-mortem human ALS and FTD tissues (Atkinson et al., 2024), which is in agreement with our previous results in TDP-43^A315T^ mice showing that alterations to TDP-43 are linked to reduced circulating levels of leptin and the dysfunction of its signalling pathways at the central nervous system (CNS) (Ferrer-Donato et al., 2021). These data are of interest because studies have suggested that higher leptin levels are critical for survival in ALS disease, as leptin levels have been found to be lower in sporadic ALS patients with rapidly progressing disease (Picher-Martel et al., 2023), which is in agreement with our previous studies in TDP-43^A315T^ mice showing the potential benefit of leptin therapy against ALS (Ferrer-Donato et al., 2022). Thus, it will be important to study the possible alterations occurring in WAT, which could perturb peripheral to CNS communication, leading to skeletal muscle degeneration in ALS. Here we aimed to better understand whether the WAT plays a critical role in the pathophysiology of ALS, as most ALS patients have hypermetabolism (Fayemendy et al., 2021) and weight loss (Dardiotis et al., 2018) prior to the onset of motor symptoms. In this context, considering the potential remodelling of WAT and its important role in whole-body homeostasis, we comprehensively characterized changes in WAT at different stages of the disease (asymptomatic, onset and end-stage) in TDP-43^A315T^ mice compared to gender and age- matched wild-type (WT) littermates, using histology, proteomics method, and molecular biology techniques.

In addition, as epidemiological evidence suggests, alterations in leptin levels are associated with changes in disease progression and survival in ALS. Plasma samples of patients with ALS were used to assess sex and BMI differences in leptin concentrations at the time of diagnoses, to determine if leptin may serve as prognostic biomarker in ALS. Exploring these links may help to better understand the mechanisms underlying the relationship between adiposity and disruption in leptin levels in ALS, because, although BMI is the most used measure of global adiposity, it does not represent adiposity in terms of regional fat distribution, which differs between sexes, races and ages.

## MATERIALS AND METHODS

### Experimental Animals

Two cohorts of sex and age-matched WT non-transgenic littermates (C57Bl/6) and TDP-43^A315T^ mice (Wegorzewska et al., 2009a) (n = 60/genotype) were used in this study to conduct histology (n = 30/genotype), proteomic and molecular biology techniques (n = 30/genotype). In both cases, the ALS-like disease was divided into three stages (n = 5/stage): asymptomatic, onset (defined as the last day of individual peak body weight before a gradual loss occurs) and the end-stage of disease (defined as when weight is 20% below the initial weight on three consecutive days), which is typically reached 2–4 weeks after symptom onset. To avoid the ambiguity associated with reported sex-related differences in mean survival time of this mouse model of ALS (Hatzipetros et al., 2014; Wegorzewska et al., 2009b), only male mice were used. Animals were euthanized independently on each stage of disease. Animals expressing the human TDP-43 (hTDP-43) transgene were confirmed via PCR according to the distributor’s protocol.

To monitor disease onset and progression, all mice were weighed and assessed three times per week until the disease onset-stage, after which they were checked daily in the morning until the disease end-stage. The maintenance and use of mice and all experimental procedures were approved by the Animal Ethics Committee of the Hospital Nacional de Parapléjicos, Toledo (Spain) (Approval No 26/OH 2018) in accordance with the Spanish Guidelines for the Care and Use of Animals for Scientific Purposes. All analyses were conducted by personnel blinded to the animal genotype.

### Perfusion and tissue collection

Animals were terminally anesthetized with sodium pentobarbitone (140 mg/kg, intraperitoneally) and transcardially perfused with 0.01 M PBS (pH 7.4). For immunohistochemistry, subcutaneous WAT (scWAT) and perigonadal WAT (pgWAT) were immediately dissected, rinsed in cold phosphate buffered saline (PBS), postfixed 70% ethanol, and stored at 4ªC until paraffin embedding using an Automatic Tissue Processor (ATP 1000, Histo-line, Italy), for further use. For molecular biology experiments, scWAT and pgWAT tissues of each animal were split into two fractions and processed independently, for real time qPCR and Western blot analysis, or proteomics methods. Samples were immediately frozen on dry ice and stored at −80 °C for later analysis.

### Immunohistochemistry (IHC)

Paraffin-embedded sections (4μm thick) of scWAT and pgWAT were deparaffinized in xylene and rehydrated through descending grades of ethanol (100%, 95%, 90%, 80% and 70%) to water. Sections were stained with hematoxylin and eosin (H&E), dehydrated through ascending grades of ethanol (75%, 95%, and 100%), cleared in xylene and finally mounted in dibutyl phthalate xylene (DPX). For analysis, digital photomicrographs of the entire scWAT and pgWAT tissue sections (20x; Leica CTR 6000, Leica, Mannheim, Germany) were used to quantify the histochemical staining in 10 random fields for each sample different regions to assess the regional heterogeneity in tissue samples. The regions were outlined using ImageJ software (Rasband, W.S., ImageJ, U. S. National Institutes of Health, Bethesda, Maryland, USA, https://imagej.net/ij/, 1997-2018). All slides were analysed using the same morphologic criteria for the quantification of crown-like structures (CLSs), defined by the clustering of macrophages (identified by each morphology) to surround a dying adipocyte, as a sign of adipose-tissue inflammation. This criterion was: presence of a ring of mononuclear cells surrounding an adipocyte vacuole. In addition, the number of blood vessels (BV) was also determined to assess vascularity as an index of angiogenesis.

Finally, by semiquantitative methods we determined inflammatory markers such as mononuclear infiltrate and fibrosis, and necrosis markers such as adipocyte normal membrane shape and tissue integrity. Semiquantitative scores were assigned by an observer based on predefined morphologic criteria: 1 (<25%), 2 (25–50%), 3 (51– 75%), and 4 (>75%). Measure 10 fields at 10x in simple scWAT and pgWAT tissues (n = 60). All analyses were conducted by personnel blinded to animal genotype.

### Total protein preparation and mass spectrometry analysis Sample preparation for library generation

Proteins were separated on precast gel, 4–20% MiniPROTEAN TGX (BioRad) and visualized by Coomassie staining. The entire gel lane was manually cut into 8 sections and subjected to in-gel tryptic digestion. The digestion was performed according to Schevchenko *et al*. (Shevchenko et al., 1996) with minor modifications: gel slices were incubated with 10mM dithiothreitol (DTT; Sigma Aldrich) in 50mM ammonium bicarbonate (99% purity; Scharlau) for 60min at 37°C and after reduction, alkylation with 55mM iodoacetamide (IAA; Sigma Aldrich) in 50mM ammonium bicarbonate was carried out for 20min at RT. Gel plugs were washed with 50mM ammonium bicarbonate in 50% methanol (gradient, HPLC grade, Scharlau), rinsed in acetonitrile (ACN, gradient, HPLC grade, Scharlau) and dried in a Speedvac. Dry gel pieces were then embedded in sequencing grade modified porcine trypsin (Promega, Madison, WI, USA) at a final concentration of 12.5ng/µL in 20 mM ammonium bicarbonate. After digestion at 37 °C overnight, peptides were extracted with 60% acetonitrile in 0.5% formic acid (FA, 99.5% purity; Sigma Aldrich) and the samples were resuspended in 10µL [98% water with 2% formic acid and 2% ACN].

### Sample preparation for SWATH analysis

Samples lysates were digested using Single-pot solid-phase-enhanced sample preparation (SP3) according to the protocol of Hughes et al. The lysates were reduced and alkylated using DTT and IAA, respectively. After reduction and alkylation, 6 µL of the prepared bead mix was added to the lysate and made up to 30 µL using H2O. Afterward, EtOH was added to a final concentration of 70% (v / v) and the samples were left stirring at 1000rpm and room temperature for 20 min. Subsequently, the beads were immobilized by incubation on a magnetic rack for 2 min. The supernatant was recovered in a new vial and the entire procedure was repeated. The pellet was rinsed with 80% (v / v) EtOH in water several times. The beads were resuspended in 300 µl of 100mM NH4HCO3 supplemented with trypsin in an enzyme to protein ratio of 1:25 (w / w). After digestion overnight at 37°C and 1000 rpm, the samples are centrifuged at 20,000 g, the supernatant is collected and acidified using 2% FA.

### LCMSMS Analysis for library, DDA and SWATH

In order to build the spectral library, the peptides extract were analysed by a shotgun approach by nanoLC-MS/MS. Samples were pooled and 3µg was separated into a Ekspert^™^ nanoLC425 (Eksigent, Dublin, CA, USA) using a C18 column (ChromXPC18, 3µm, 120Å 0.075 x 150 mm, Eksigent) at a flow rate of 300nL/min in combination with a precolumn (NanoLC Trap ChromXP C18, 3µm 120Å, Eksigent) at a flow rate of 5µL/min. The buffers being used were: A = 0.1%FA 2%ACN and B = 98% ACN in water with 0.1% FA. Peptide were desalted for 3 min with 0.1%FA/2% ACN on the precolumn, followed by a separation for 85min using gradient from 5% to 30% solvent B, 30%-95% for 0.1min, and finally 95%B for 5min. Column was then regenerated with 5%B for 10 additional minutes. Peptides eluted were directly injected into a hybrid quadrupole-TOF mass spectrometer TripleTOF^®^ 6600+ (Sciex, Redwood City, CA, USA). Sample was ionized in a source type Optiflow < 1µL Nano applying 3.0kV to the spray emitter at 200°C. Analysis was carried out in a data-dependent positive ion mode (DDA). Survey MS1 scans were acquired 350-1400 m/z for 250 ms. The TripleTOF was operated in SWATH mode, in which a 50 ms TOF MS scan from 350–1400 m/z was performed, followed by 50 ms product ion scans from 100–1500 m/z on the 70 variable windows from 350 to 1400 Da (2.20 sec/cycle). The individual SWATH injections were randomized.

### Protein Data Analysis

Peptide and protein identifications were performed using ProteinPilot^TM^ Software V 5.0 (Sciex) and the Paragon algorithm (Shilov et al., 2007). Each MS/MS spectrum was searched against the uniprot- proteome_MusMusculus_2021_04 database, with the fixed modification of carbamidomethyl -labelled cysteine parameter enabled. Other parameters such as the tryptic cleavage specificity, the precursor ion mass accuracy and the fragment ion mass accuracy, are TripleTOF^®^ 6600plus built in functions of the ProteinPilot software. SWATH Acquisition MicroApp v.2.0 was used for building a peptide spectral library containing the peptide identified in the database search with confidence score above 95%. SWATH Acquisition MicroApp was used for extracting the ion chromatogram traces from the SWATH raw files and using the previously generated spectral library, and the following parameters: 20 peptides/protein; 6 fragment ions/peptide; extraction windows of 5 min and 25 ppm; peptide FDR of 1% and confidence score threshold of 95%. Normalisation of the protein abundance signal as measured by SWATH was carried out using MarkerView (v1.2.1, Sciex).

### RNA Isolation and qPCR analysis

Total RNA was isolated from WAT using the RNeasy Mini Kit (Qiagen), according to the manufacturer’s instructions. Complementary DNA (cDNA) (0.5 µg of total RNA) synthesis and the relative quantification of *TDP-43*, *UCP-1*, *C/EBPβ* and *PPARγ* were performed as described previously (Fernandez et al., 2009). The 18S rRNA was used as a control to normalize gene expression (Fernandez et al., 2010). The reactions were run on an CFX96 Real-Time System instrument and software (CFX Manager 3.0) (BioRad) according to the manufacturer’s protocol. Primers were designed using NCBI/Primer-BLAST software (Table 1). Relative quantification for each gene was performed by the ΔΔCt method (Livak & Schmittgen, 2001).

### Protein extraction and western-blot analysis

Proteins from WAT were extracted using RIPA buffer (Sigma Aldrich) containing a cocktail of protease inhibitors (Roche) as described previously (Ferrer-Donato et al., 2021). Denatured protein samples (20 µg) from each group were electrophoresed into Bolt® Bis–Tris Plus gels (Invitrogen), transferred to PVDF membranes (BioRad) and incubated with rabbit anti-TDP-43 (1:1000; Proteintech) overnight. Subsequently, anti-rabbit horseradish peroxidase (HRP)-conjugated secondary antibody (Vector Laboratories) was used as described previously (10.3390/ijms221910305). Mouse anti-actin (1:1000; Cell Signalling) was used as a loading control and band intensity was measured as the integrated intensity using ImageJ software (v1.4; NIH). All data were normalized to control values on each membrane.

### Human plasma samples

All procedures performed in studies involving human samples (plasma) were in accordance with the Ethics Committee (783/23/98) of the University CEU-San Pablo, Madrid, Spain. Human plasma samples were provided by the Biobank HUB-ICO-IDIBELL, integrated in the ISCIII Biobanks and Biomodels Platform and they were processed following standard operating procedures with the appropriate approval of the Ethics and Scientic Committees.

Patients were eligible for inclusion if they had diagnosis of ALS based on Gold Coast criteria (Shefner et al., 2020). Controls were also provided by the Biobank HUB-ICO-IDIBELL. Samples were collected at the hospital using Lithium heparin collecting tubes at the time of diagnosis. Samples were centrifuged at 10,000 RPM for 10 min. The supernatant was collected and frozen at -80 degrees Celsius.

ALS patients were classified by sex, BMI and survival. Patients were classified as nonobese (BMI, <25) and obese (BMI, ≥25). BMI was calculated at the time of diagnosis. Patients were classified as slow progressors when survival was higher than 5 years, as normal progressors when survival was between 3 and 5 years, and as fast progressors when survival was less than 3 years. All the human samples included were of patients who were already deceased at the time of the study.

### Measurement of plasma leptin by ELISA

ELISAs were performed as suggested by the manufacturer’s protocol. Leptin plasma levels were measured using a Human Leptin ELISA Kit PicoKine® (Boster Biological Technology) with samples diluted 1:20 for males and 1:10 for females. Each patient’s samples were processed in duplicate. The limit of detection was 62.5 pg/mL, and the within-assay and between-assay coefficient of variability (CVs) were 7.8% and 6.5%, respectively.

### Statistical Analysis

Statistical analyses were performed using GraphPad Prism software v10.2.0. Normality of datasets was assessed by Kolmogorov-Smirnov Test. Outliers were removed with ROUT method with Q = 1 %. For IHC Mann-Whitney test was used. For molecular biology analysis, two-way ANOVA was used followed by Dunett’s post hoc test to compare all groups with control WT asymptomatic mice, while Tukey’s post hoc test was used for multiple comparisons between all groups. To compare within the same group, t-test test was used. For ELISA, two-way ANOVA was used followed by Tukey’s post hoc test for multiple comparisons between all groups. Mann-Whitney test was used to compare within the same group. A Spearman correlation coefficient (rho) was employed to assess the correlation between quantitative variables, with significance set at a p-value of ≤ 0.05 (n=78). This analysis was performed with SPSS Statistics. Values were reported as means ± standard error of the mean (SEM). For all comparisons, significant results were taken when p value *<* 0.05.

## RESULTS

### Histological examination of WAT reveals significant alterations in TDP-43^A315T^ mice

Given the limited understanding of how WAT is altered during the progression of ALS, we aimed to address this gap by performing a comprehensive histological characterization of scWAT and pgWAT tissues across the three stages of the disease -asymptomatic, onset and end-stage- in TDP-43^A315T^ mice, compared to age- matched WT littermates, using H&E staining analysis. We first evaluated CLSs and the number of BV in histological sections (Fig. 1; Suppl. Fig. 1 and 2), and IHC analysis demonstrated marked differences in the number of both parameters in scWAT and pgWAT, respectively, during the clinical course of the disease in TDP-43^A315T^ mice compared with WT samples (Fig. 1B; Suppl. Fig. 1B and 2B). Although the number of CLSs was similar regardless of the location of the adipocytes (scWAT *vs.* pgWAT) (Fig. 1B), Mann-Whitney test demonstrated a statistically significant increase in the number of CLSs in both scWAT and pgWAT tissues during the asymptomatic stage in TDP-43^A315T^ mice (Fig. 1B). The number of BV were also similar regardless of the location of the adipocytes (scWAT *vs.* pgWAT) (Fig. 1C), however, Mann-Whitney test demonstrated a statistically significant decrease in the number of BV in both scWAT and pgWAT in TDP-43^A315T^ mice compared to WT mice, suggesting an alteration in the vascularity of the WAT of TDP-43^A315T^ mice (Fig. 1C). In contrast, at the onset stage of disease, no statistically differences were found in both parameters analysed between TDP-43^A315T^ *vs.* WT samples (Suppl. Fig. 1B-C), however, Mann-Whitney test demonstrated a statistically significant increase in the number of CLSs and BV in both scWAT and pgWAT tissues at the end- stage of the disease in TDP-43^A315T^ mice compared to age-matched WT littermates (Suppl. Fig. 2B-C).

**Fig. 1.**
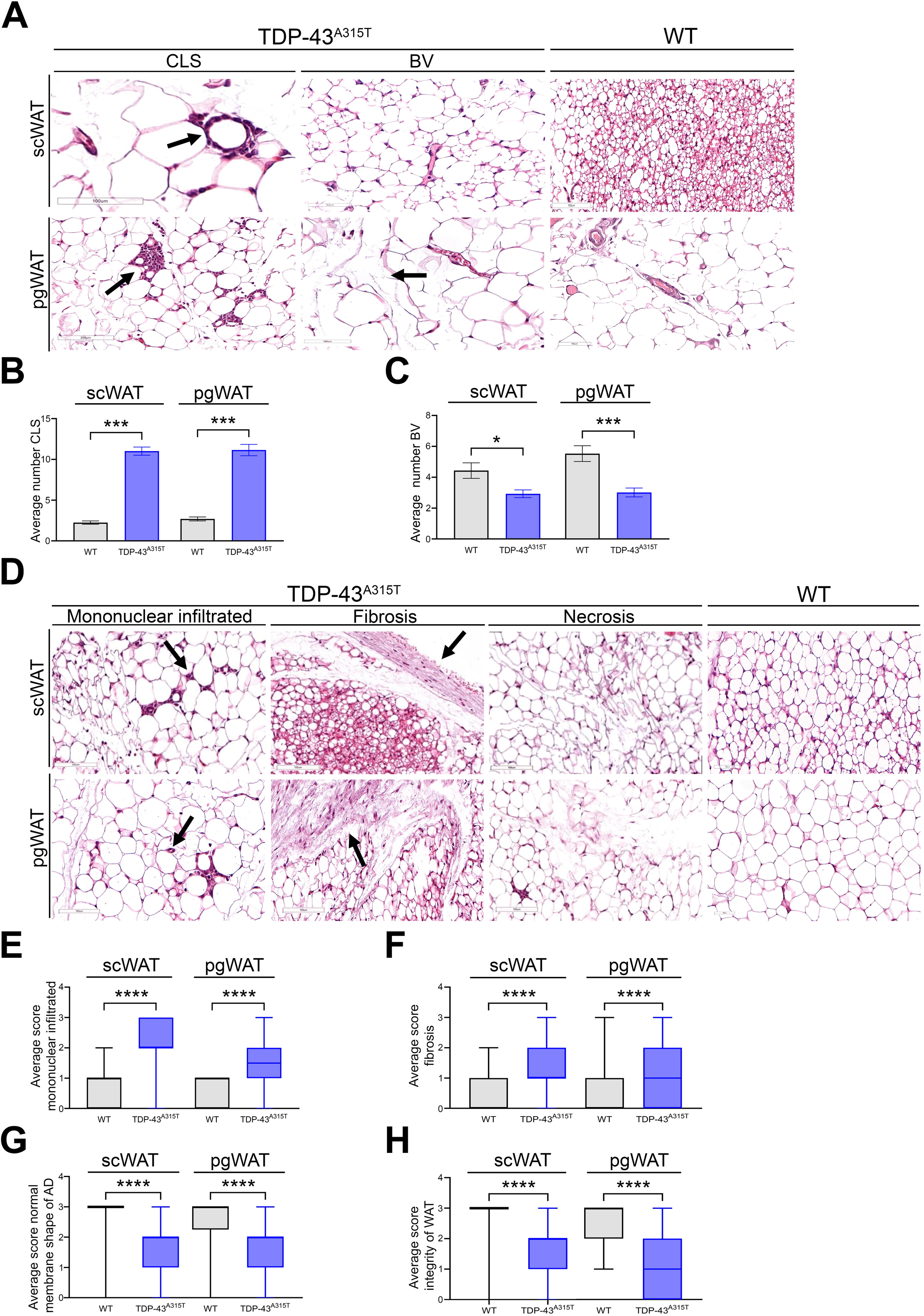
Hematoxylin and eosin stained scWAT and pgWAT in asymptomatic stage of male TDP-43^A315T^ mice. (A) Representative images of CLSs and BV (section bar = 100 µm). (B) Semi-quantitative analysis of CLSs and (C) BV. (D) Representative images of mononuclear infiltrated, fibrosis and necrosis markers. (E) Semi-quantitative analysis of mononuclear infiltrated, (F) fibrosis, (G) normal membranes shape of AD and (H) integrity of WAT. Abbreviations: scWAT, subcutaneous adipose tissue; pgWAT, perigonadal adipose tissue, CLS, crown-like structures; BV, blood vessels; AD, adipocyte; WAT, white adipose tissue.

**Fig. 2.**
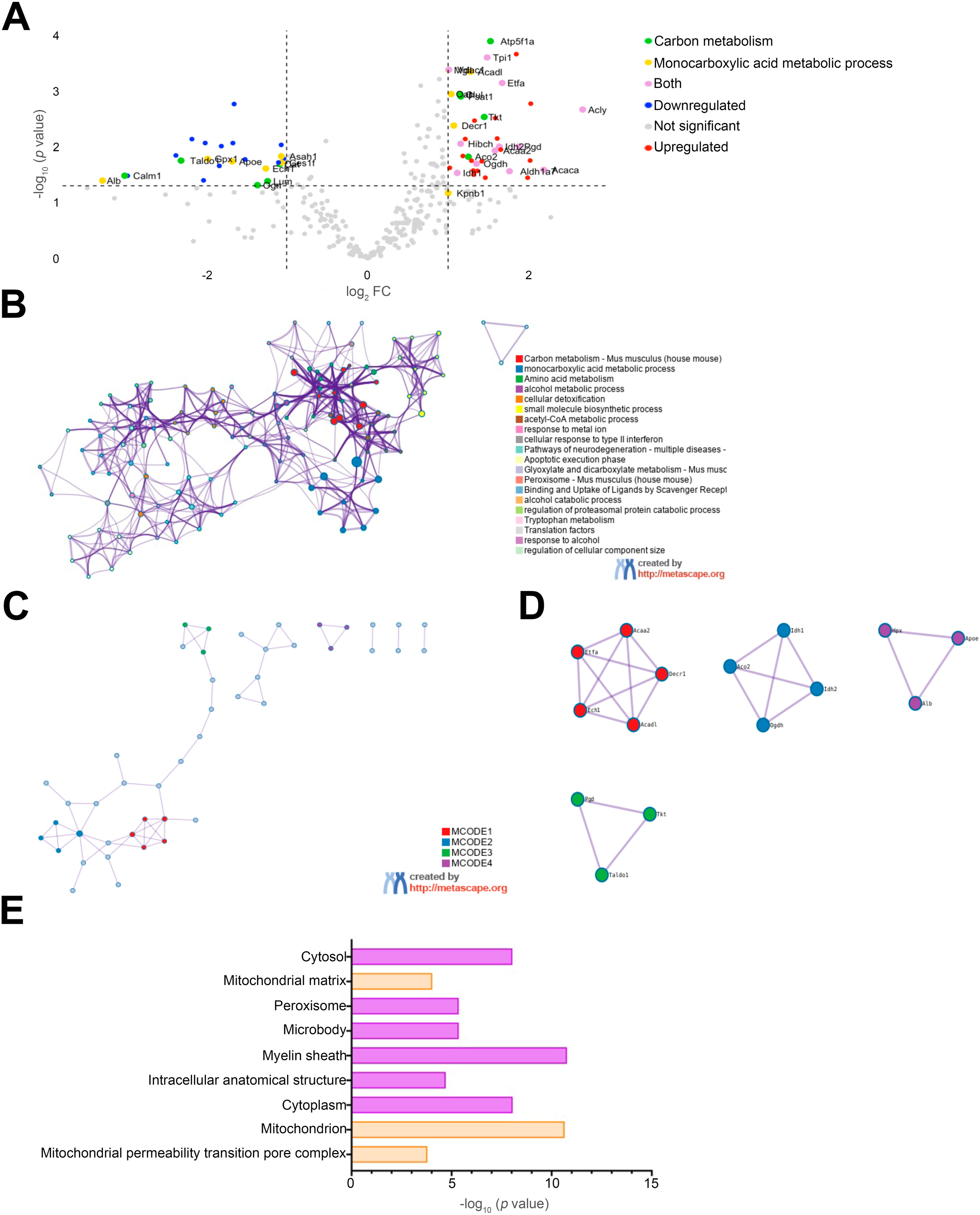
Proteomic analysis of WAT at asymptomatic stage in TDP-43^A315T^ mice. (A) Volcano plot. Each point represents an individual protein. (B) Enrichment map of representative enriched ontology terms. (C) PPI network colored by MCODE clusters colored by Metascape. (D) PPI network of the separated MCODE clusters from panel C. (E) Cellular component enrichment of DEPs generated with Gene Ontology Software. Abbreviations: WAT, white adipose tissue; PPI, protein-protein interaction; DEPs, proteins differentially expressed.

To gain deeper histological insight into the pathological modifications occurring in WAT during the progression of ALS, we further evaluated mononuclear infiltrate, fibrosis and necrosis in both scWAT and pgWAT tissues at the three stages of the disease in TDP-43^A315T^ mice *vs.* WT samples (Fig. 1D-H; Suppl. Fig. 1 and 2D-H, respectively). IHC analysis demonstrated a statistically significant increase on inflammatory marker of mononuclear infiltrate at the asymptomatic stage (Fig. 1E) and during the symptomatic stage of disease (Suppl. Fig.1 and 2E, respectively) in TDP-43^A315T^ mice compared to WT mice. Mann-Whitney test demonstrated a statistically significant increase on the fibrosis in both scWAT and pgWAT tissues TDP-43^A315T^ mice compared to WT controls (Fig. 1F; Suppl. Fig. 1 and 2F, respectively). Regarding adipocyte shape and tissue integrity there was a significant decrease in both scWAT and pgWAT tissues in TDP-43^A315T^ mice (Fig. 1G-H; Suppl. Fig. 1 and 2G-H, respectively).

### Proteomics analysis of WAT from TDP-43^A315T^ mice highlighted the presence of mitochondrial alterations prior to the onset of motor symptoms

To further elucidate the molecular alterations occurring in WAT of TDP-43^A315T^ mice, we conducted a proteomic analysis of pgWAT. We focused our analysis on pgWAT, as in this type of WAT fibrosis and inflammation is increasingly appreciated as a major player in adipose tissue dysfunction, and pgWAT had substantially higher pro-inflammatory characteristics than scWAT. In this context, using a significant threshold of p<0.05, proteins having FC>2 were determined as up-regulated, whereas the ones with FC<0.5 were down- regulated.

Focusing on the asymptomatic stage of the disease, a total of 1528 proteins were detected as proteins differentially expressed (DEPs) in TDP-43^A315T^ mice compared to age-matched WT littermates. Of these DEPs, 38 showed to be upregulated, 24 downregulated, and 1466 showed no statistically significant differences (Fig. 2A). Enrichment and ontology analyses were performed in Metascape, considering upregulated and downregulated DEPs. Enriched ontology cluster network showed significant enrichment of terms mainly related to two categories: carbon metabolism and monocarboxylic acid metabolic process (Fig. 2C). MCODE clustering analysis recognized 4 different clusters in the protein-protein interaction (PPI) network (Fig. 2D-E; Table 2). MCODE1 represented ontology terms related to mitochondrial fatty acid beta-oxidation and respiratory electron transport. MCODE2 represented ontology terms related with mitochondrial oxidoreductases and ligase and MCODE4 with oxidoreductase and transferases. Though, MCODE3 represents ontology terms related with heme-degradation, scavenging of heme from plasma, lipid-transport. Cellular enrichment analysis showed DEPs grouped together in mitochondria, peroxisome and cytosol (Fig. 2F).

Furthermore, focusing on the end-stage of the disease, 121 DEPs were identified from a total of 1569 proteins in TDP-43^A315T^ mice compared to age-matched WT littermates. Among these DEPs, 75 showed to be upregulated, 46 downregulated and 1448 showed no statistically significant differences (Fig. 3A). Upregulated and downregulated DEPs were considered for the enrichment and ontology analyses performed in Metascape. Enriched ontology cluster network showed significant enrichment of terms mainly related to three categories: cellular catabolic process, terpenoid metabolic process and metabolism of xenobiotics by cytochrome p450 (Fig. 3C). MCODE clustering analysis recognized 2 different clusters in the PPI network (Fig. 3D-E; Table 2). MCODE1 represented ontology terms related to lyases, transferases, oxidoreductases, proteins transport, ribonucleoproteins and RNA and DNA binding. MCODE2 represented ontology terms related to transferases, monooxygenases and oxidoreductase. Cellular enrichment analysis showed DEPs grouped together in cytoplasm, endoplasmic reticulum and mitochondrion, among others (Fig. 3F).

**Fig. 3.**
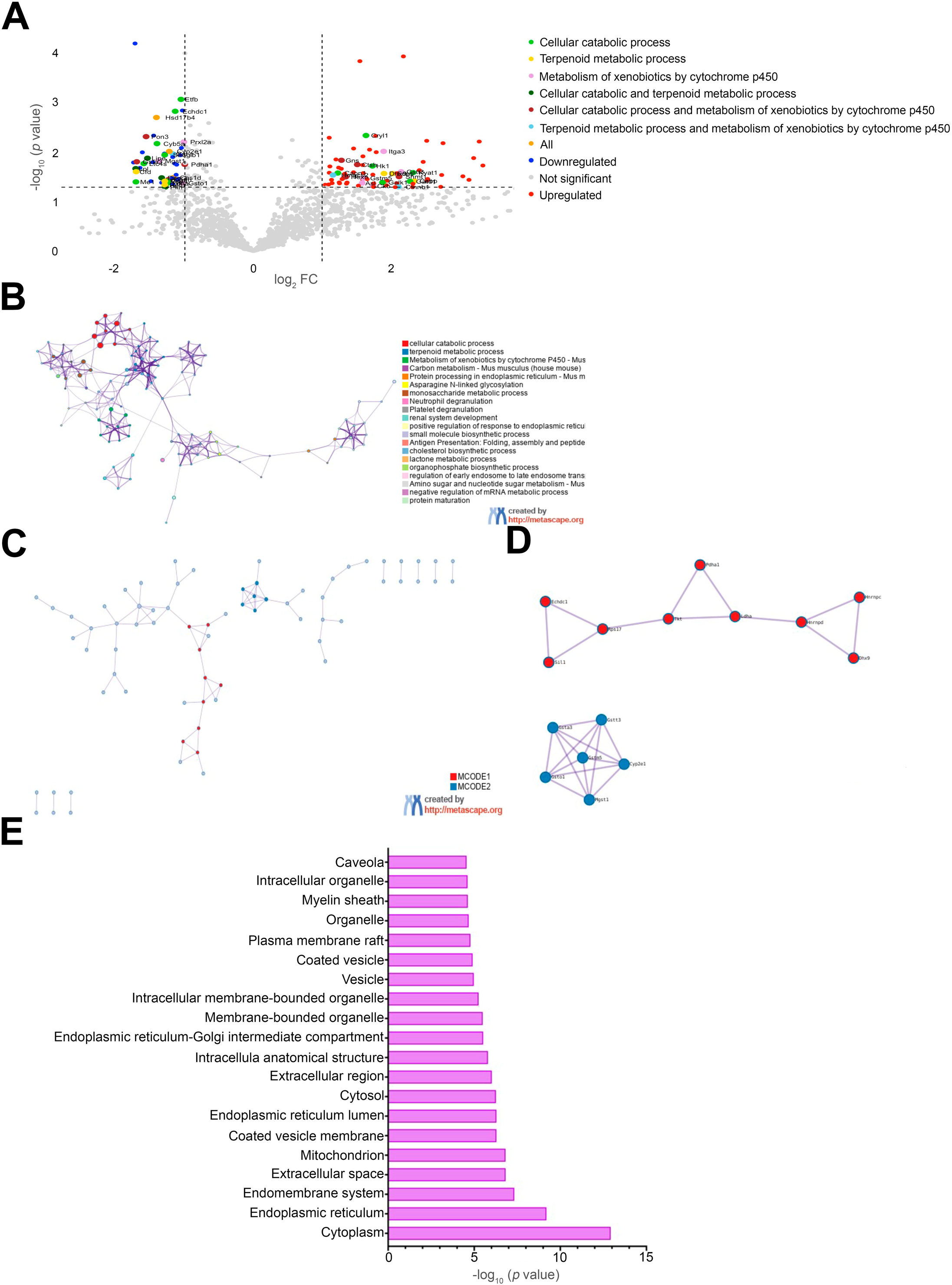
Proteomic analysis of WAT at end-stage in TDP-43^A315T^ mice. (A) Volcano plot. Each point represents an individual protein. (B) Enrichment map of representative enriched ontology terms. (C) PPI network colored by MCODE clusters colored by Metascape. (D) PPI network of the separated MCODE clusters from panel C. (E) Cellular component enrichment of DEPs generated with Gene Ontology Software. Abbreviations: WAT, white adipose tissue; PPI, protein-protein interaction; DEPs, proteins differentially expressed.

Table 2 reports the list of MCODE cluster recognized by the integrated Metascape analysis of proteomic data showed in Fig. 2D-E and Fig. 3D-E. For each cluster, the top 3 enriched terms are reported, relatively to the lowest *p*-value.

### WAT of TDP-43^A315T^ mice display altered *PPARγ* mRNA expression levels concomitantly with an increase on the protein levels of TDP-43

Proteomic analysis highlighted several alterations in pgWAT at the asymptomatic stage in TDP-43^A315T^ mice, highlighting the presence of mitochondrial alterations, which could potentially impact both cellular differentiation and metabolism. Thus, to better characterize such defects we performed RT-qPCR in the pgWAT of TDP-43^A315T^ mice over the time course of the disease. RT-qPCR analysis demonstrated no statistically significant differences in the mRNA expression levels of *C/EBPβ* and *PPARγ* between TDP- 43^A315T^ and WT mice at either of the time-points analysed (Fig. 4A-B). In addition, RT-qPCR analysis detected no *Ucp1* mRNA in the pgWAT. We then sought to better characterize the patterns of TDP-43 transgene expression in pgWAT of TDP-43^A315T^ mice compared to age-matched WT littermates. In concordance with the pathological modifications occurring in pgWAT prior to the onset of motor symptoms in TDP-43^A315T^ mice, endogenous *TDP-43* mRNA levels (mTDP-43) were significantly upregulated in TDP-43^A315T^ mice compared to WT mice during the asymptomatic stage (*p* = 0.005; Fig. 4C), while no statistically significant differences were found between TDP-43^A315T^ and WT mice at both onset and end-stage. Interestingly, western blot analysis showed that TDP-43 protein levels in pgWAT, probed with a polyclonal antibody that recognizes both human and mouse TDP-43, were increased in TDP-43^A315T^ mice compared to WT mice at either of the time-points analysed (Fig. 4D-F; Suppl. Fig.3).

**Fig. 4.**
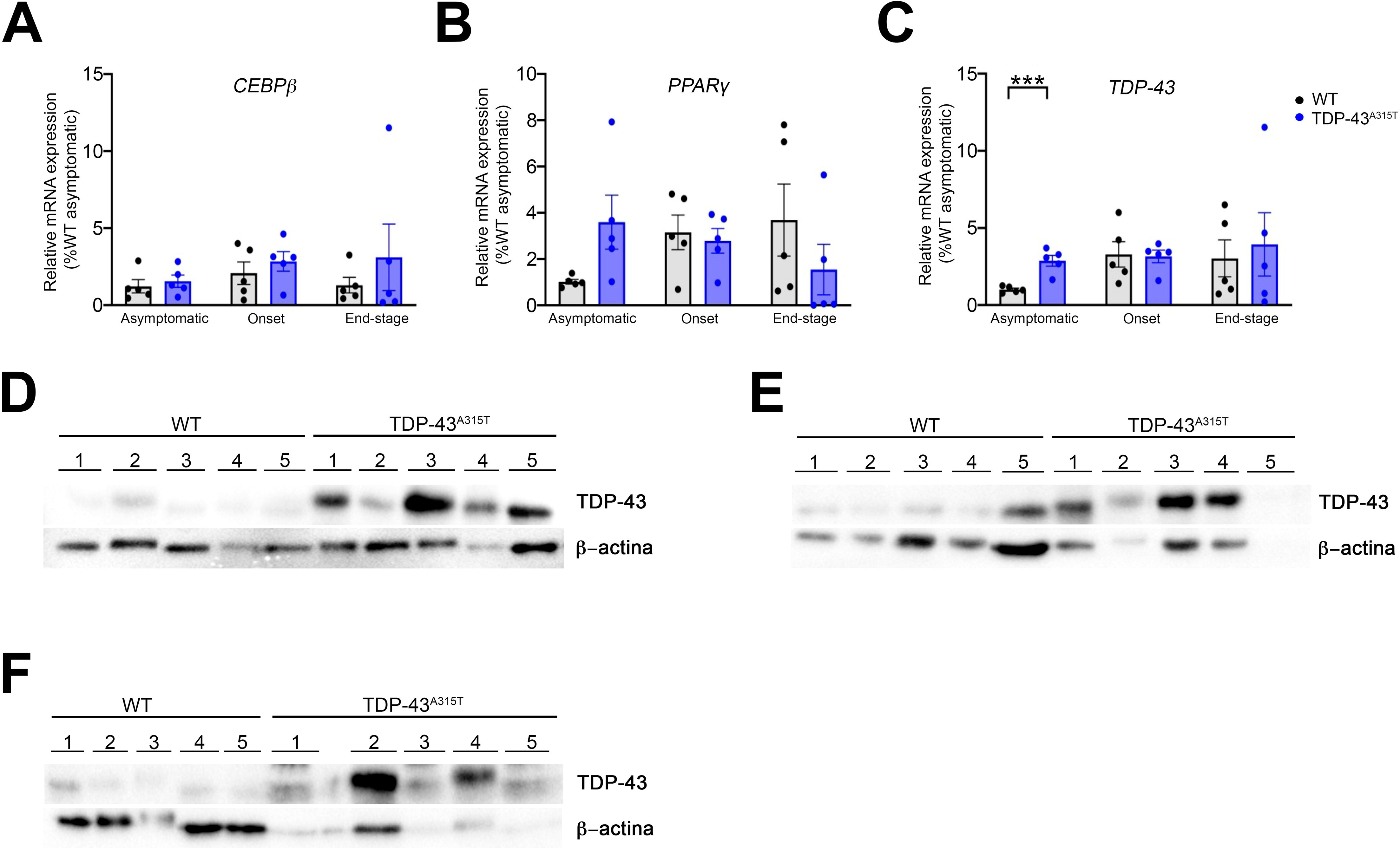
Alterations in *C/EBPβ, PPARγ* and *TDP-43* in WAT of TDP-43^A315T^ mice. (A) *C/EBPβ* (B) *PPARγ* and (C) *TDP-43* mRNA expression. Transcripts were assessed by RT-qPCR in TDP-43^A315T^ mice compared to age-matched WT littermates at asymptomatic, onset and end-stage of disease. Values are expressed as the mean ± SEM for the different groups. TDP-43 protein in WAT extracts of TDP-43^A315T^ mice compared to age- matched WT littermates at asymptomatic (D), onset (E) and end-stage of disease (F). \**p <* 0.05, \*\**p <* 0.01, \*\*\**p <* 0.001, \*\*\*\**p <* 0.0001. Abbreviations: C/EBPβ, CCAAT Enhancer Binding Protein Beta; PPARγ, Peroxisome proliferator-activated receptor gamma; TDP-43, TAR DNA-binding protein 43; WAT, white adipose tissue; WT, wild-type.

### Distinct plasma leptin profile in obese men ALS cases with rapidly progressive disease

It is unclear whether changes in WAT precede the first signs of ALS, and whether these alterations are manifested systemically through fluctuations in plasma leptin levels, which are known to be influenced by sex and age. Thus, to better understand the mechanisms regulating WAT disruption and the sexual dimorphism in circulating leptin levels in ALS patients. In total, we measured leptin levels in plasma samples of 62 ALS patients and 16 age-matched controls, with 37 men and 41 women of a wide age (30 - 87 yr) and BMI (19.06 – 33.90 kg/m2) range. No statistically significant difference between patients and control groups was found regarding age (*p* = 0.5749) nor BMI (*p* = 0.6940) (Table 3).

ELISA analysis showed that protein levels of leptin in plasma are increased in ALS patients compared to controls, while it is not significant (Fig. 5A). Sorting data by sex, protein levels of leptin were significantly increased in plasma of men ALS patients compared to men controls, as well as in women ALS patients when compared to men ALS patients and to men controls (Fig. 5B, *p* = 0.019, *p* < 0.001 and *p* < 0.001, respectively), while no statistically significant differences were found in women ALS patients compared to women controls. However, when stratifying patients by survival rate, it is observed that plasmatic leptin levels decreased significantly in men ALS patients that have had a fast disease progression when compared to women ALS patients with normal disease progression (Fig. 5C, *p* = 0.040). On the contrary, leptin levels in women are similar across the different types of progression, being significantly increased in ALS fast progression women when compared with ALS fast progression men (Fig. 5C, *p* < 0.001). Interestingly, when stratifying patients by BMI, for overweight ALS patients, the decrease in plasma leptin levels was significant in men ALS patients with fast disease progression when compared to women ALS patients with normal disease progression (Fig. 5D, *p* = 0.006). Moreover, leptin levels of men ALS patients with normal disease progression were also significantly decreased when compared to women ALS patients with normal progression (Fig. 5D, *p* = 0.013). For overweight women ALS patients, levels decreased in both slow and fast progression when compared to levels in women ALS patients with normal progression, although not significance was reached (Fig. 5D).

**Fig. 5.**
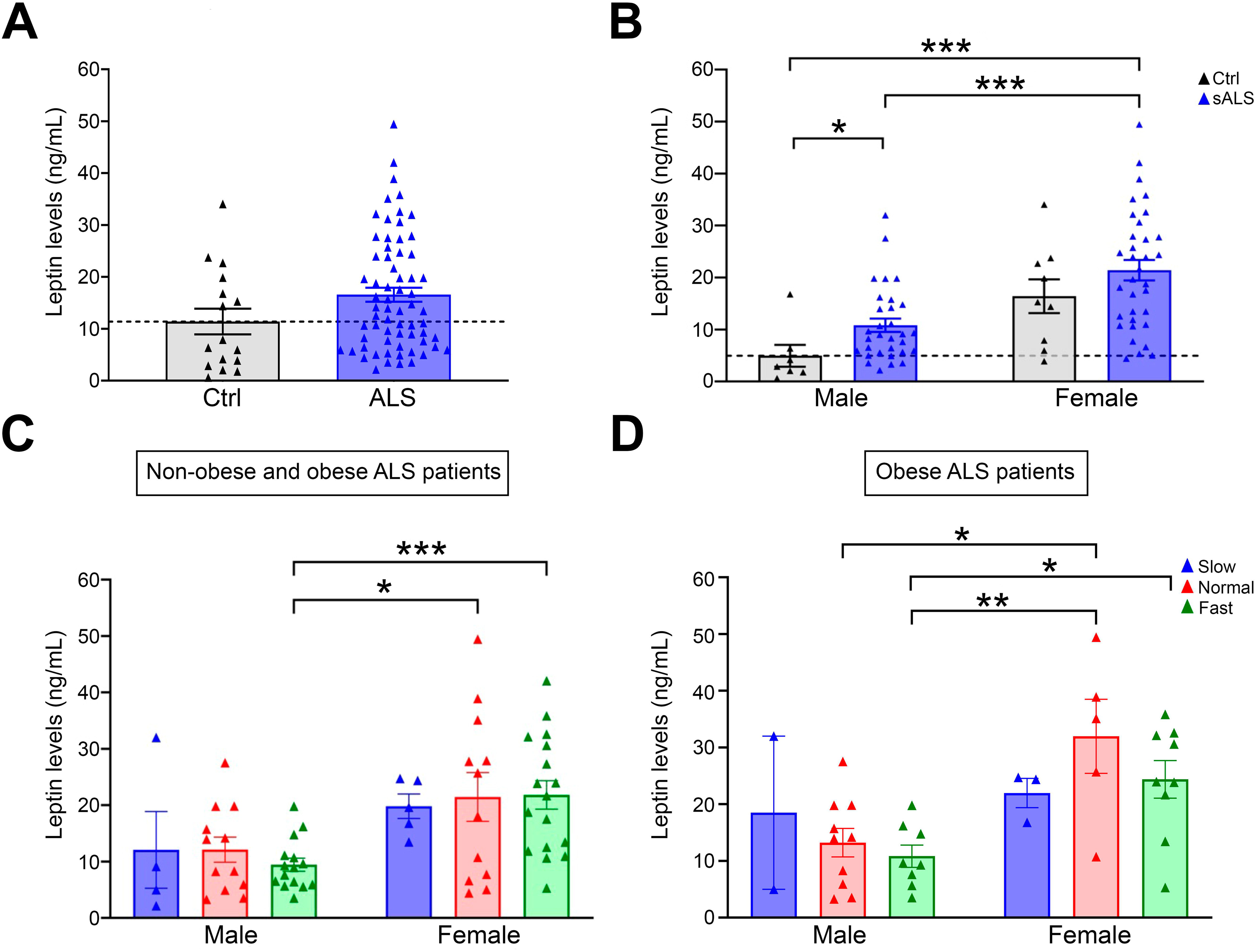
Analysis of plasma leptin levels in ALS patients. (A) Concentration of leptin levels at diagnosis measured by ELISA in healthy controls and ALS subgrouped by sex. (B) Concentration of leptin levels at diagnosis in ALS and controls subgrouped by sex and disease progression: slow, normal and fast progression patients. (C) Concentration of leptin levels at diagnosis in ALS and controls subgrouped by sex and rate of progression in overweight patients. \**p <* 0.05, \*\**p <* 0.01, \*\*\**p <* 0.001, \*\*\*\**p <* 0.0001. Abbreviations: ALS, sporadic amyotrophic lateral sclerosis.

Furthermore, bivariate correlations performed are showed in Table 4 and both sex and BMI exhibit a monotonic relationship with leptin levels (*p* < 0.001, r = 0.510 and *p* < 0.001, r = 0.434, respectively). The correlation of leptin plasma levels with survival rate, as well as the correlation between the qualitative variables, were also evaluated, but no significant differences were found.

## DISCUSSION

Although metabolic dysfunctions of the CNS in ALS have been widely studied (Rosina et al., 2025), metabolic alterations observed in ALS patients and animal models have not been well investigated in peripheral organs such as WAT, which play a key role in the endocrine control of energy homeostasis. The current study aimed to better understand whether the WAT plays a critical role in the pathophysiology of ALS, which is of interest as determining how restoring adipose tissue plasticity may contribute significantly to mitigate the hypermetabolism observed in patients with ALS. In this context, we identify for the first time evidence of a pathological dysfunction in WAT prior to the symptomatic stage of the disease in TDP-43^A315T^ mice, providing novel insights about the pathways that could link dysregulating systemic energy homeostasis to the progression of ALS. Additionally, an important finding of our study was that circulating leptin levels at the time of diagnosis, were lower in the plasma of men with ALS who were overweight or obese and had rapidly progressive ALS, emphasizing the importance of considering sex-specific approaches to guide the development of effective clinical therapies.

At present, there is increasing interest in the use of hypercaloric diets (e.g. high-fat diet; HFD), as gaining weight and, subsequently fat mass has been associated with better survival in ALS (Heritier et al., 2015). Both preclinical and human research demonstrate a disease-modifying effect of nutritional state in ALS (Ngo, Shyuan T. et al., 2017). However, although epidemiological data suggest that supplementation with HFD may reduce the risk of developing ALS, as it provides positive survival outcomes (Morozova et al., 2008; Ngo, Shyuan T. et al., 2017; Okamoto et al., 2007; Veldink et al., 2007), the exact molecular mechanisms behind these effects remain elusive. Here, we report a significant pathological dysfunction of WAT prior to the symptomatic stage of the disease in TDP-43^A315T^ mice. We observed a significant increase in the number of CLSs in both scWAT and pgWAT tissues, concomitantly to a decrease in the number of BV, suggesting an alteration in the vascularity of the WAT prior to the onset of motor symptoms in TDP-43^A315T^ mice, which might negatively affect the mechanisms governing the expansion of this tissue. Our results also confirm stage- dependent alterations on inflammatory marker of mononuclear infiltrate in TDP-43^A315T^ mice. These observations are of interest because inflamed adipocytes are a key feature of this condition as they secrete, both locally and systemically, proinflammatory cytokines, such as tumor necrosis factor-alpha (TNFα), which in turn disrupt the normal function of adipose tissue itself and compromises the inefficient expandability of WAT. Indeed, we have previously reported in a more refined model of TDP-43 proteinopathy, the rNLS8 mice (Walker et al., 2015), that the ability of a short-term HFD therapy to improve prognosis in ALS mice is not a simple question of gaining weight, which produce a systemic low-grade inflammation itself, it may depend on the capacity of WAT to respond to a caloric excess by its healthy expansion (Romero-Muñoz L et al., 2024). Indeed, an impaired WAT remodelling results in alterations in lipid stored, leading to metabolic derangements, such as altering adipokines production and release (Martinez-Sanchez, 2020), which supports previous data from our group showing a significantly downregulation of peripheral leptin levels from the pre-onset stage of disease in TDP-43^A315T^ mice (Ferrer-Donato et al., 2021), thereby influencing metabolic circuits involved in energy intake regulation.

Consistently with earlier findings, we also provided data showing significant changes in the proteomic profile of WAT in TDP-43^A315T^ mice, highlighting alterations such as mitochondrial dysfunction, which can alter the cellular homeostasis of WAT and the adipocytes (Chen et al., 2023), concomitantly with an increase on the protein levels of TDP-43 in different phases of the disease compared to age-matched WT littermates. This observation is of interest because the evidence supports that TDP-43 is a powerful regulator of body fat composition (Stallings et al., 2013) by mechanisms involving the transcriptional regulation of genes that impaired leptin signalling (Dokas et al., 2016), thereby contributing to leptin insensitivity. Thus, it is conceivable that increased protein levels of TDP-43 in WAT disrupt the production and release of leptin by the adipocytes and contribute to a dysfunction of metabolic homeostasis in the CNS in TDP-43^A315T^ mice, as we previously reported alterations in leptin signalling in the spinal cord and the hypothalamus compared to WT controls (Ferrer-Donato et al., 2021), which is in agreement with novel data from our group showing disruption to leptin signalling in pathology-rich brain regions of postmortem human tissue of ALS (Atkinson et al., 2024). However, future experiments should try to corroborate this hypothesis.

In patients, adipose tissue distribution is altered and has been correlated with functional status and survival (Lindauer et al., 2013). Sex differences in disease-related endocrine dysfunction have also been confirmed (Grassano et al., 2024), which may explain, at least in part, the higher susceptibility to ALS in men than women (Handley et al., 2022). However, although the mechanisms driving this dimorphic incidence are still largely unknown, in accordance with previous published research conducted, both in humans (Hellstrom et al., 2000; Konukoglu et al., 2000) and mouse model of ALS (Picher-Martel et al., 2023), we observed marked sex differences in the plasma levels of leptin early on the symptoms in ALS patients. Accordingly, ELISA analysis demonstrated that leptin levels were significantly increased in the plasma of men, but not in women with ALS, compared to control subjects at the time of diagnosis, which is of interest because it has been established that leptin levels in females are generally higher than in male due to their higher percentage of body fat and their different hormonal profile (Hellstrom et al., 2000). However, when stratifying patients by progression of the disease, it is observed that plasmatic leptin levels decreased significantly in men ALS patients that have had a fast disease progression compared to women with ALS. Interestingly, when stratifying patients by BMI, for overweight patients, the decrease in plasma leptin levels was significant in men with fast disease progression compared to women with ALS and with both normal and fast disease progression, respectively. These data suggest that women may be more protected than men against ALS, due to at least two different potential mechanisms, which are not mutually exclusive. One possible mechanism could be the neuroprotective effect of female hormones in the early stages of ALS disease. Indeed, published data showed how sex dichotomy in ALS is only present in the pre-menopausal aged female population, indicating a potential protective role of circulating estrogen (Manjaly et al., 2010). Alternatively, sex-related differences in endocrine dysfunction associated with the disease could suggest that pathological dysfunction of WAT is less severe in women than in men ALS patients. Indeed, our unpublished new experimental data reveal a significant delay in the pathological disturbances observed in both scWAT and pgWAT tissues of female TDP-43^A315T^ mice compared to male ALS mice and WT samples, respectively, which is consistent with reported experimental data in SOD1^G93A^ mice (Picher-Martel et al., 2023). It is also noteworthy that leptin levels are higher in overweight or obese women than in overweight or obese men (Konukoglu et al., 2000), consistent with a state of relative leptin resistance, and reflecting differences in body fat composition. Thus, this data might indicate the differences in the physiological response of the hypothalamus to overcome WAT atrophy in ALS, highlighting both leptin and WAT as a putative target organ or to have a prognostic/diagnostic significance.

In conclusion, we demonstrate an impairment of WAT during the manifestation of ALS phenotype, which undergoes significant alterations that could potentially impact on the normal physiology of the adipocytes. We showed a significant increase in the number of CLSs, a characteristic histopathology feature of inflamed WAT (Murano et al., 2013), an alteration in its vascularity, which promotes adipocyte dysfunction and induces oxidative stress, hypoxia and inflammation (AlZaim et al., 2023), a stage-dependent alteration on inflammatory marker of mononuclear infiltrate, and significant changes in the proteomic profile, highlighting mitochondrial alterations, which could significantly disrupt leptin levels (Cavaliere et al., 2023), during the clinical course of disease in TDP-43^A315T^ mice. These alterations in WAT could facilitate a powerful crosstalk between metabolically active organs (e.g. liver, brain, heart and kidneys), modulating energy homeostasis in TDP-43^A315T^ mice. Indeed, although it remains speculative that such alterations could be due to WAT developmental defects, and although it is important to bear in mind when interpreting our experimental results in TDP-43 mice, they may not be transferable to the clinical practitioner. Further research in ALS patients is crucial to deciphering the role of WAT and understanding the specific functions of leptin levels in the metabolic dysfunction caused by pathological TDP-43, the cause of which remains elusive.

## List of abbreviations

ACN: acetonitrile
AgRP: agoute-related peptide
ALS: amyotrophic lateral sclerosis
ATP: Automatic Tissue Processor
BAT: brown adipose tissue
BMI: body mass index
BV: blood vessels
cDNA: complementary DNA
CLSs: crown-like structures
CNS: central nervous system
CVs: coefficient of variability
DDA: data-dependent positive ion mode
DEPs: proteins differentially expressed
DPX: dibutyl phthalate xylene
DTT: dithiothreitol
FA: formic acid
FTD: frontotemporal dementia
H&E: hematoxylin and eosin
HFD: high-fat diet
HRP: horseradish peroxidase
hTDP-43: human TDP-43
IAA: iodoacetamide
IHC: Immunohistochemistry
mTDP-43: endogenous TDP-43
PBS: phosphate buffered saline
pgWAT: perigonadal WAT
POMC: antagonist pro-opiomelanocortin
PPI: protein-protein interaction
scWAT: subcutaneous WAT
SEM: standard error of the mean
SP3: single-pot, solid-phase-enhanced
TDP-43: TAR DNA binding protein
TNFα: Tumor Necrosis Factor alpha
UFMN: Functional Motorneuron Unit
WAT: white adipose tissue
WT: wild-type

## Declarations

### Ethics approval and consent to participate

All animal procedures were performed in accordance with the Animal Ethics Committee of the Hospital Nacional de Parapléjicos (Approval No 36OH/2019) (Spain) in accordance with the European Communities Council Directive (86/609/EEC) for the Care and Use of Animals for Scientific Purposes.

Human samples described in this manuscript were collected under protocols that were reviewed and approved by the Functional Unit of Amyotrophic Lateral Sclerosis (UFELA), Service of Neurology, Bellvitge University Hospital, Hospitalet de Llobregat, (Spain). Subjects underwent informed consent prior to participating in research studies.

### Consent for publication

All authors have read and agreed to the published version of the manuscript.

### Availability of data and materials

The datasets used and/or analysed during the current study available from the corresponding author on reasonable request.

### Authors’ contributions

C-BC, A-FD collected the tissues; C-BC, E-DM performed histological data analysis; G-BG performed proteomic and provided expertise in data analysis; R-DM, MP provided clinical expertise in data analysis and patient information; C-BC, C-FM performed statistical analysis and prepared the figures. C-FM conceived and designed the study. C-FM drafted the manuscript. All authors critically reviewed the manuscript for intellectual content. All authors have read and agreed to the published version of the manuscript.

## Acknowledgments

The authors would like to thank the Proteomics Unit of the Hospital Nacional de Parapléjicos for their invaluable technical and analytical assistance in the use of proteomic technology, and the Surgery Unit of the Hospital Nacional de Parapléjicos, Toledo (Spain) for their excellent technical support. In addition, the authors would like to thank patients and Biobank HUB-ICO-IDIBELL (PT20/00171) integrated in the ISCIII Biobanks and Biomodels Platform and Xarxa Banc de Tumors de Catalunya (XBTC) for their collaboration and provide human plasma samples. Finally, Cristina Benito-Casado is supported by a PhD Fellowship from the Consejería de Educación, Ciencia y Universidades Comunidad de Madrid (PIPF-2023/SAL-GL-29613).

---

**Suppl. Fig. 1.**
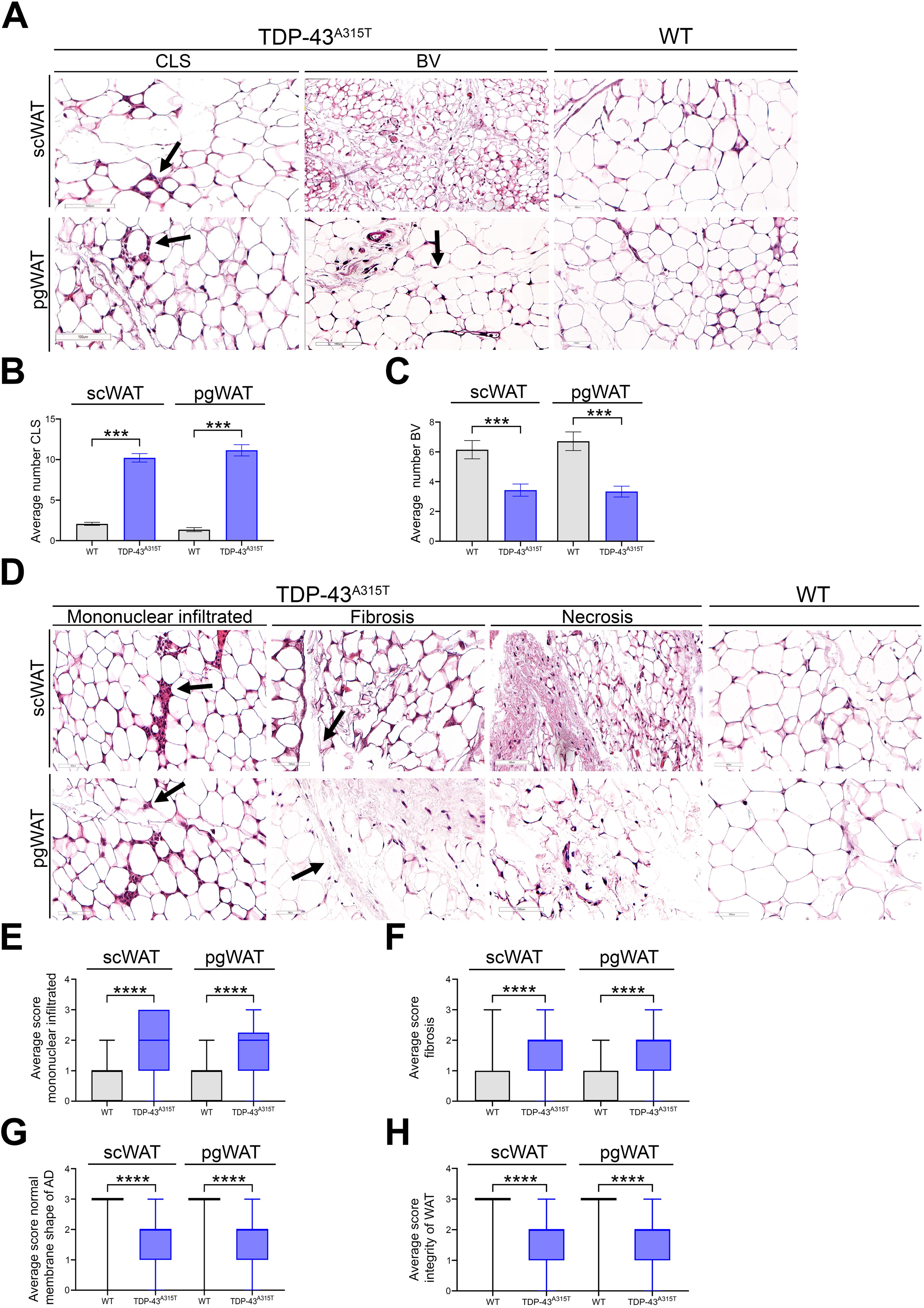
Hematoxylin and eosin stained scWAT and pgWAT in onset stage of TDP-43^A315T^ mice. (A) Representative images of CLSs and BV (section bar = 100 µm). (B) Semi-quantitative analysis of CLSs and (C) BV. (D) Representative images of mononuclear infiltrated, fibrosis and necrosis markers. (E) Semi- quantitative analysis of mononuclear infiltrated, (F) fibrosis, (G) normal membranes shape of AD and (H) integrity of WAT. Abbreviations: scWAT, subcutaneous adipose tissue; pgWAT, perigonadal adipose tissue, CLS, crown-like structures; BV, blood vessels; AD, adipocyte; WAT, white adipose tissue.

**Suppl. Fig. 2.**
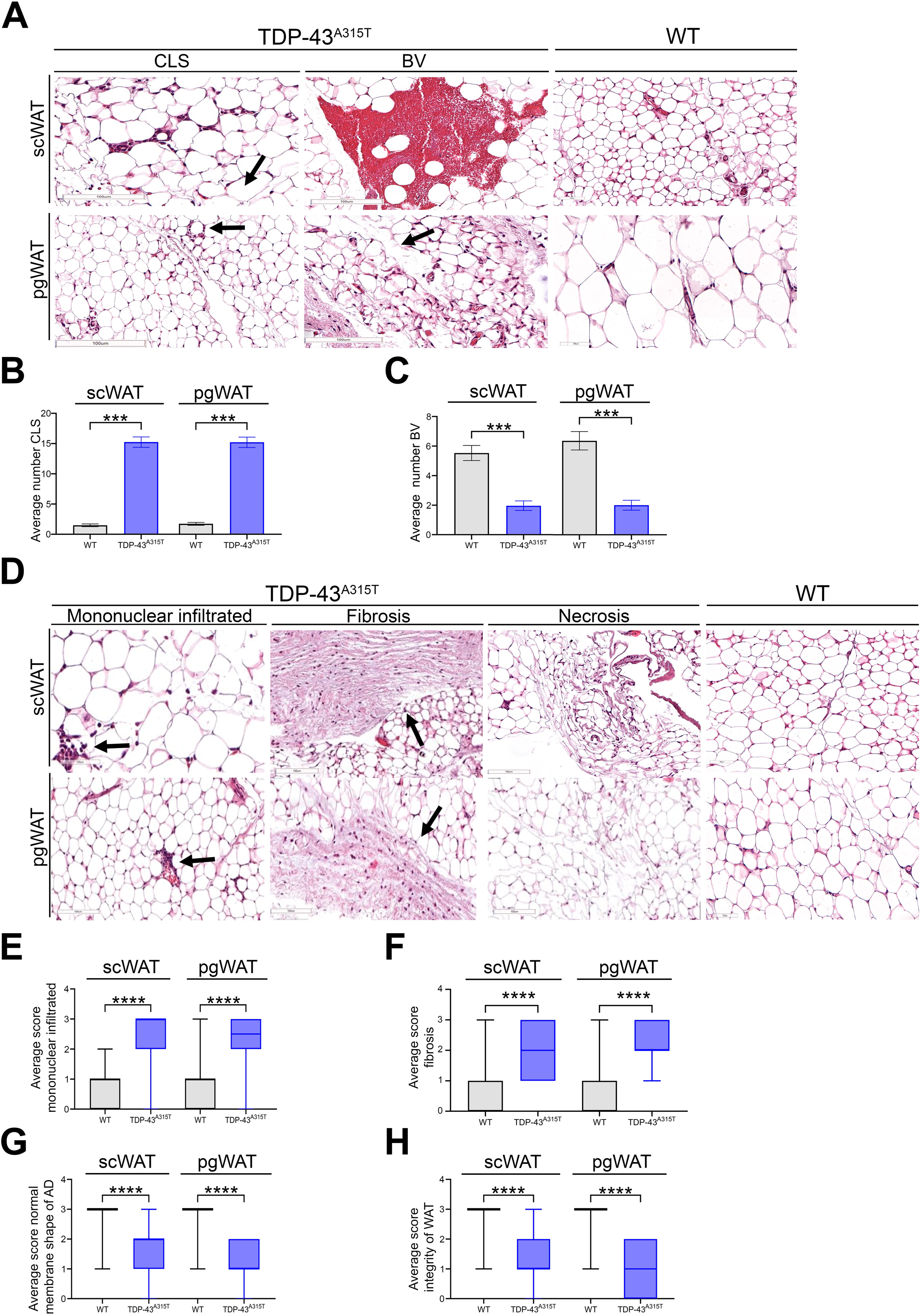
Hematoxylin and eosin stained scWAT and pgWAT in end-stage of TDP^A315T^ mice. (A) Representative images of CLS and BV (section bar = 100 µm). (B) Semi-quantitative analysis of CLS and (C) BV. (D) Representative images of mononuclear infiltrated, fibrosis and necrosis markers. (E) Semi- quantitative analysis of mononuclear infiltrated, (F) fibrosis, (G) normal membranes shape of AD and (H) integrity of WAT. Abbreviations: scWAT, subcutaneous adipose tissue; pgWAT, perigonadal adipose tissue, CLS, crown-like structures; BV, blood vessels; AD, adipocyte; WAT, white adipose tissue.

**Suppl. Fig. 3.**
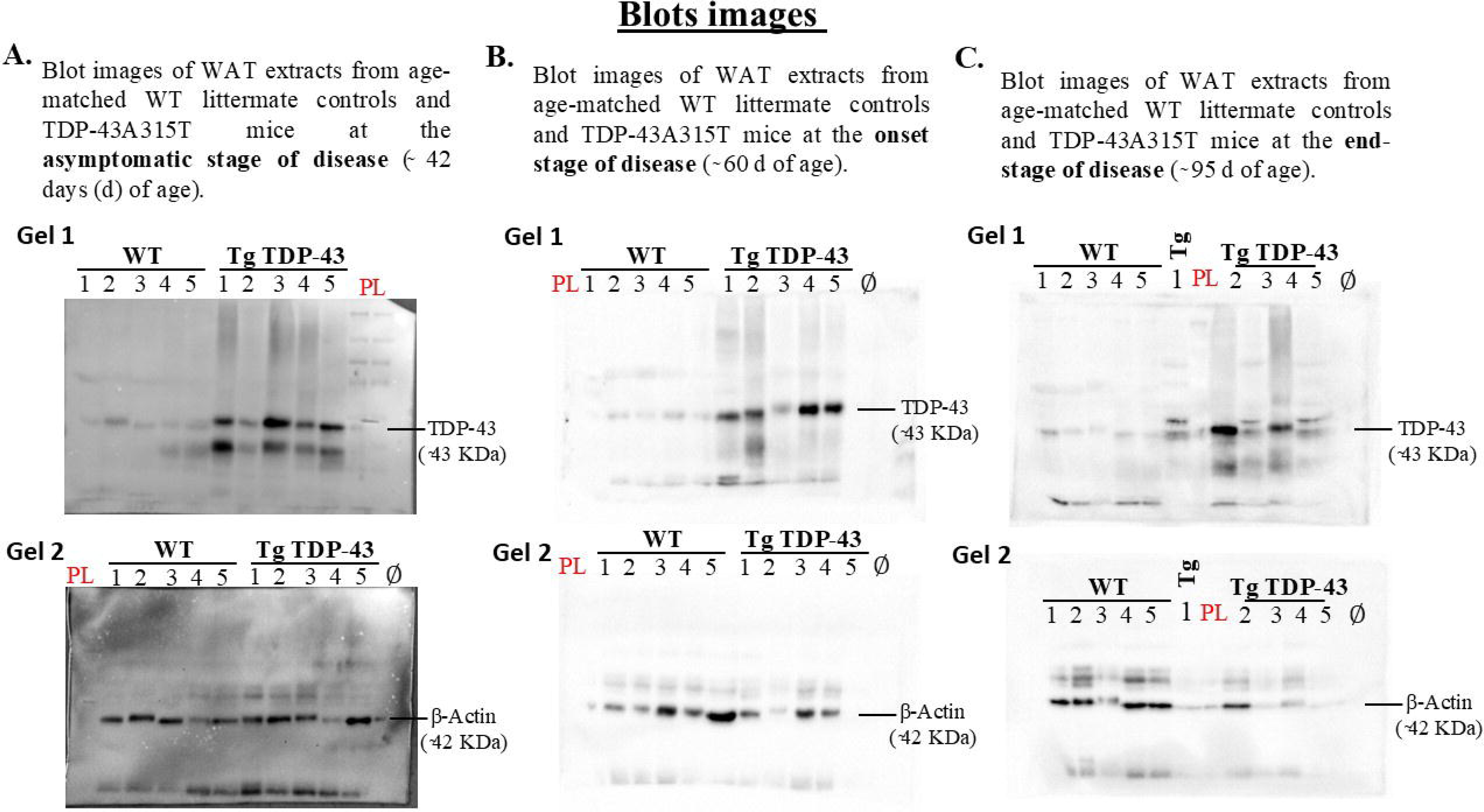
Blot images of WAT extracts at different stages of the disease (asymptomatic, onset and end-stage) labelling TDP-43 and β-Actin in TDP-43^A315T^ mice compared to gender and age-matched wild-type (WT) littermates. (A) Blot images of WAT at the asymptomatic stage of the disease. (B) Blot images of WAT at the onset stage of the disease. (C) Blot images of WAT at the end-stage of the disease. Abbreviations: WAT, adipose tissue; PL, protein ladder.

